# A proteomic feasibility study connecting metabolic and synaptic pathway alterations in serum and extracellular vesicles to characterize treatment-resistant depression

**DOI:** 10.64898/2026.08.19.745678

**Authors:** Olivia B. Ramsay, Sean A. Burnap, Matthew F. Dobbs, Weston B. Struwe, Scott J. Russo, James W. Murrough, Carol V. Robinson, Tarick J. El-Baba

**Affiliations:** Kavli Institute for Nanoscience Discovery, University of Oxford, OX1 3QU, UK; Department of Oncology, University of Oxford, OX1 3TA, UK; Department of Biochemistry, University of Oxford, OX1 3QU, UK; The Dennis S. Charney, MD, Depression and Anxiety Discovery Center, Department of Psychiatry, Icahn School of Medicine of Mount Sinai, New York, NY, USA; Nash Family Department of Neuroscience, Icahn School of Medicine at Mount Sinai, New York, NY, USA; Brain and Body Research Center, Icahn School of Medicine at Mount Sinai, New York, NY, USA; Friedman Brain Institute, Icahn School of Medicine at Mount Sinai, New York, NY, USA; VISN 2 Mental Illness Research, Education, and Clinical Center (MIRECC), James J. Peters VA Medical Center, Bronx NY; Department of Chemistry, University of Oxford, OX1 3TA, UK

## Abstract

Treatment-resistant depression (TRD) remains a major clinical challenge, yet the biological processes distinguishing TRD from non-treatment-resistant depression (nTRD) are incompletely defined. While circulating serum proteomes reflect broad systemic alterations associated with depression, extracellular vesicles (EVs) could provide a more selective representation of intercellular signaling relevant to treatment resistance. Here, we carried out a pilot study to evaluate the extent that parallel proteomic profiling of serum and serum-derived EVs could distinguish healthy controls (CON), nTRD, and TRD individuals. In this exploratory and hypothesis-generating study, serum proteomes exhibited robust global differences between depression groups and controls, largely reflecting shared systemic biology across nTRD and TRD. In contrast, EV proteomes showed limited global separation but revealed subtype-associated pathway differences. Relative to controls, nTRD EVs were enriched for immune and inflammatory pathways. By contrast, TRD EVs were characterized by enrichment of mitochondrial metabolism, oxidative phosphorylation, translational initiation, and MYC-regulated pathways, together with depletion of synaptic signalling, membrane trafficking, and cytoskeletal pathways. Comparative analysis of pathways significant in both contrasts revealed that these bioenergetic and translational signatures were selectively amplified in TRD relative to nTRD. Our exploratory analyses identified that the circulating EV cargo may reflect a treatment-resistance-specific reorganization of biological pathways not apparent in bulk serum proteomics. This study highlights parallel serum and EV proteomics as a complementary approach for molecular stratification in antidepressant resistance.

## Introduction

Major depressive disorder (MDD) affects hundreds of millions of people worldwide, and is ranked among the leading causes of years lived with disability^1^. Despite the wide availability of typical monoamine-based antidepressants, MDD broadly encompasses individuals with treatment-resistant depression (TRD) and non-TRD (nTRD). Approximately 30 to 40% of MDD patients fail to display adequate clinical improvement,^2^ and are deemed TRD.^3^ The TRD subgroup carries disproportionate morbidity, suicidality, and healthcare impact^4–6^. Yet the biological processes that distinguish TRD from nTRD (*viz.* treatment-responsive depression) remain poorly characterized. Thus, TRD continues to be defined retrospectively by clinical non-response rather than by any underlying molecular signature.

Peripheral blood-based molecular profiling has provided important biological insights into maladaptive brain processes present in psychiatric^7–10^ and central nervous system (CNS)^11–13^ disorders. Whole-blood transcriptomic profiling has identified immune, inflammatory, synaptic, metabolic, and stress-related gene expression differences between TRD, nTRD, and healthy controls, and has linked interferon signalling pathways and transcriptional changes to ketamine treatment response^14^. Proteomic studies of peripheral fluids from MDD cohorts have similarly implicated inflammatory and metabolic dysregulation in depression^9^. However, identifying proteomic signatures that can distinguish treatment-resistant from treatment-responsive depression, thereby aid in explaining antidepressant failure, remains an unmet need.

Recent peripheral blood proteomic studies using protein-targeted O-link technologies have identified alterations in inflammatory, metabolic, and complement-related proteins in MDD^15^. Notably, overlap between MDD-associated immune signatures and those of inflammatory skin conditions has suggested avenues for drug repurposing. However, despite tremendous potential for utilizing focused proteomic studies in psychiatry, consistent molecular signatures that distinguish TRD and nTRD remain elusive. Advances in data-independent acquisition (DIA) mass spectrometry approaches, such as DIA-parallel accumulation, serial fragmentation (DIA-PASEF), now allow for substantially improved proteomic depth, enabling comprehensive coverage of complex biofluids with minimal missing data^16,17^. Yet despite these methodological advances, deep unbiased proteomic profiling of serum has not been applied to MDD, and whether such an approach can reveal molecular signatures distinguishing TRD from nTRD remains unknown.

Extracellular vesicles (EVs) are membrane-bound nanoparticles readily found in circulation and offer a conceptually distinct and complementary window into disease biology^18–20^. EVs are released by most cell types and carry protein, lipid, and nucleic acid cargo reflecting the physiological state of their cell of origin. Unlike bulk serum proteins, EV cargo is shaped by regulated packaging mechanisms, offering a more selective representation of disease-relevant intercellular signalling. Critically, EVs can traverse the blood–brain barrier, raising the possibility that circulating EVs carry biologically informative cargo of CNS origin^20^. This has motivated substantial interest in exploiting EVs as peripheral liquid biopsies for neuropsychiatric disorders. Accordingly altered EV-associated proteins, miRNAs, and lipids have been reported across schizophrenia,^21^ bipolar disorder,^22^ and MDD^23,24^. However, progress has been hampered by methodological limitations, particularly the widespread reliance on L1CAM immunocapture to enrich for putative neuron-derived EVs^25^; in both serum and cerebrospinal fluid. L1CAM has been shown to be present predominantly as a soluble, cleaved or alternatively spliced protein^26^. Thus, establishing strategies for *bona fide* enrichment of CNS-derived EVs using surface molecules is an active area of research.^27^

Recent methodological advances in unbiased EV enrichment now permit high-depth proteomic profiling, tractable from small biofluid volumes^28^. The Mag-Net approach employs a magnetic bead-based EV enrichment strategy that sieves membrane-bound particles from plasma using strong anion exchange chemistry, enabling identification of thousands of proteins beyond the dynamic range of unfractionated biofluid analysis, including proteins with restricted CNS expression. Whether such unbiased EV proteomics can improve the molecular stratification of TRD relative to nTRD remains to be explored.

In the present study, we performed parallel DIA-PASEF proteomic profiling of serum and serum-derived EVs from a pilot cohort of healthy controls and individuals with MDD that are nTRD or TRD. Following adjustment for demographic covariates, we applied differential abundance and pathway-level analyses to characterise compartment- and subtype-specific proteomic alterations. We show that bulk serum proteomes capture robust but largely shared systemic alterations across depressive subtypes. By contrast, EV proteomes reveal a treatment-resistance-specific reorganisation of biological pathways not apparent in bulk serum, and are characterised by amplified bioenergetic and translational pathways and relative depletion of synaptic and membrane trafficking signatures. Our findings suggest that compartment-selective proteomic profiling of circulating EVs may offer an ideal approach for molecular stratification of antidepressant treatment resistance.

## Materials and Methods

### Patient characteristics and blood draws

MDD was assessed with the Structured Clinical Interview for the Diagnosis and Statistical Manual of Mental Disorders – fifth edition.^29^ MDD symptom severity was evaluated using Quick Inventory of Depressive Symptomatology-Self Report (QIDS-SR).^30^ Healthy control (CON) patients did not meet the criteria for MDD diagnosis and were not evaluated for symptom severity. MDD participants were stratified TRD (n=7) or nTRD (n = 8). Here, TRD was defined as lack of response to two or more established antidepressant treatments in the current episode at the time of screening. On the day of the blood draw, patients were fasted for >6 h. Blood was drawn into Vacutainer Gold Top 5 mL Silica Gel tubes (BD, #365968). Blood was then allowed to clot for >30 min, centrifuged at 1,300 g for 15 min at 4 °C, aliquoted, and stored at -80 °C until use. The median age across all participants was 32 years (range 22–63), with 52.2% female participants. Cohort characteristics are detailed in **Table 1**.

**Table 1.** Sociodemographic and clinical characteristics of study participants. CON, healthy control; nTRD, non-treatment-resistant depression; TRD, treatment-resistant depression; MADRS, Montgomery–Åsberg Depression Rating Scale.

| Characteristic | CON (n=8) | nTRD (n=8) | TRD (n=7) |
| --- | --- | --- | --- |
| Age, years (median [range]) | 32 [25–63] | 35 [26–52] | 32 [22–45] |
| Female sex, n (%) | 4 (50%) | 4 (50%) | 4 (57%) |

### Serum proteomics sample preparation

Serum samples (10 μL) were clarified by centrifugation at 20,000 ×g (4 °C, 30 min) and subjected to depletion of the top 14 high-abundance proteins using a Pierce Top-14 depletion column according to the manufacturer’s instructions. Proteins were denatured in 6 M urea, reduced with 10 mM TCEP, alkylated with 5 mM iodoacetamide, and digested overnight with trypsin at 37 °C. Tryptic peptides were desalted and resuspended at approximately 0.05 μg μL^-1^ in 98:2 water:acetonitrile containing 0.1% formic acid prior to liquid chromatography-tandem mass spectrometry (LC-MS/MS) analysis.

### Extracellular vesicle isolation and proteomics

EVs were isolated from 100 μL of clarified serum using a MagNet-based automated purification workflow on a KingFisher Flex system. EV proteins were processed by the same reduction, alkylation, and tryptic digestion protocol described for total serum proteomics. Tryptic peptides were desalted and resuspended as described above prior to LC-MS/MS analysis.

### LC-MS/MS and data analysis

Purified peptides (50 ng; 1 μL injection) were separated by nanoflow reversed-phase liquid chromatography (nanoElute 2, Bruker Daltonics) using a 30 min gradient. Peptides were resolved on a 25 cm × 75 μm analytical column packed with 1.7 µm C18 particles (120 Å pore size; Aurora Ultimate CSI, IonOpticks) at 50 °C and a flow rate of 250 nL min^-^¹. Mobile phase A consisted of 0.1% formic acid in water and mobile phase B of 0.1% formic acid in acetonitrile. The gradient was 2–23% B over 18 min, 23–35% B over 4 min, and 35–90% B over 4 min, followed by a 4 min wash at 90% B.

Eluting peptides were analyzed on a TIMS quadrupole time-of-flight mass spectrometer (timsTOF Ultra, Bruker Daltonics) via electrospray ionization. Source conditions were 4.5 kV capillary voltage, 3.0 L min^-^¹ dry gas flow, and 180 °C dry gas temperature. Data was acquired in dia-PASEF mode over an m/z range of 400–1,000 and an ion mobility range of 1/K₀ = 0.64–1.45 Vs cm^-^². The TIMS analyzer operated at 100% duty cycle with 100 ms accumulation and 100 ms ramp times (cycle time 0.96 s). Collision energy was applied in a mobility-dependent manner, increasing linearly from 20 to 59 eV.

DIA-PASEF raw data files (.d) were processed using DIA-NN^31^ (v2.0) with a human proteome spectral library derived from the UniProt human proteome database. False discovery rate was controlled at 1% at both precursor and protein levels. Cross run normalisation was enabled using default DIA-NN settings and label-free quantification (LFQ) values were extracted and used for all downstream analyses. Protein intensities were log_2_-transformed prior to filtering and those detected in fewer than three biological replicates per group were excluded from analysis. Missing values present in ≥60% of samples within a group were imputed using a Perseus-style downshifted normal distribution approach (width 0.3, downshift 1.8 standard deviations). Remaining sporadic missing values were handled by applying a pseudocount prior to log transformation (log_2_[LFQ + 1]).

### Statistical and pathway analyses

Prior to all differential analyses, log_2_-transformed protein intensities were adjusted for age and sex using a linear model of the form log__2__(Abundance□) = β_0_□+□β_1_·Age□□+ □β_2_·Sex□□+ ε□, and the residuals were used for downstream analyses. Principal component analysis (PCA) was performed in Python (version 3.13.0) on covariate-adjusted protein matrices. Differential abundance testing, also done in Python, was carried out on covariate-adjusted residuals, generating moderated t-statistics for each contrast (nTRD vs CON; TRD vs CON). A nominal p-value<0.05 was used as significance threshold given the exploratory nature of this study.

Gene ontology (GO) enrichment analysis was performed using PANTHER^32^ Overrepresentation Test (Released 20240807) on the Gene Ontology Consortium website (geneontology.org). For differential abundance contrasts, GO enrichment was performed separately on downregulated and upregulated proteins from nTRD vs CON and TRD vs CON comparisons. Where appropriate, GO enrichment was performed on the combined to 20 proteins with high PC1 loading (from both TRD and nTRD groups). proteins were tested against the Human reference proteome using a Fisher’s exact test with false discovery rate (FDR) correction, with GO Biological Process complete annotations (GO Ontology database, Released 2025-10-05). Enriched terms were filtered to FDR < 0.05 and redundant terms were manually collapsed to retain the broadest representative term within each biological theme.

Gene set enrichment analysis (GSEA) was performed in R (version 4.5.2) using limma^33^ and fgsea packages within the Bioconductor environment, with default settings unless otherwise noted. Gene sets were drawn from the Hallmark (H) and curated Reactome (C2:CP:REACTOME) collections from MSigDB^34^ (v2026.1.Hs). Enrichment scores were calculated using 10,000 permutations. Only gene sets containing between 10 and 50% of the detected protein universe were retained. Enrichment significance was defined at FDR < 0.05. For pathways significant in both contrasts, a differential enrichment metric (ΔNES = NES_TRD–CON_ – NES_nTRD–CON_) was calculated to quantify relative pathway dysregulation associated with treatment resistance. ΔNES values are descriptive and not subjected to downstream statistical testing.

## Results

### Cohort characteristics and serum proteomic data quality

The purpose of this study was to evaluate the extent that paired serum and EV proteomes can be used to distinguish treatment groups within a psychiatric cohort. The study therefore is comprised of a modest 23 participants (CON n = 8, nTRD n = 8, TRD n = 7); we focused on balancing age (median age 32 years) and sex representation (52.2% female; **Table 1**; **Fig. 1a**). Following top-14 depletion, DIA-PASEF acquisition and quality filtering, 1,800 proteins were quantified across serum samples. The ranked abundance distribution confirmed a broad dynamic range (log_10_ LFQ 2–7; **Fig. 1b**), and per-sample protein abundance distributions were highly consistent (**Fig. 1c**). Pairwise correlations were high (Pearson R ∼0.95; **Fig. 1d**), with within-group correlations consistently exceeding between-group comparisons (**Fig. 1e**), indicating high group-associated biological variation relative to technical variability in this exploratory cohort.

**Figure 1.**
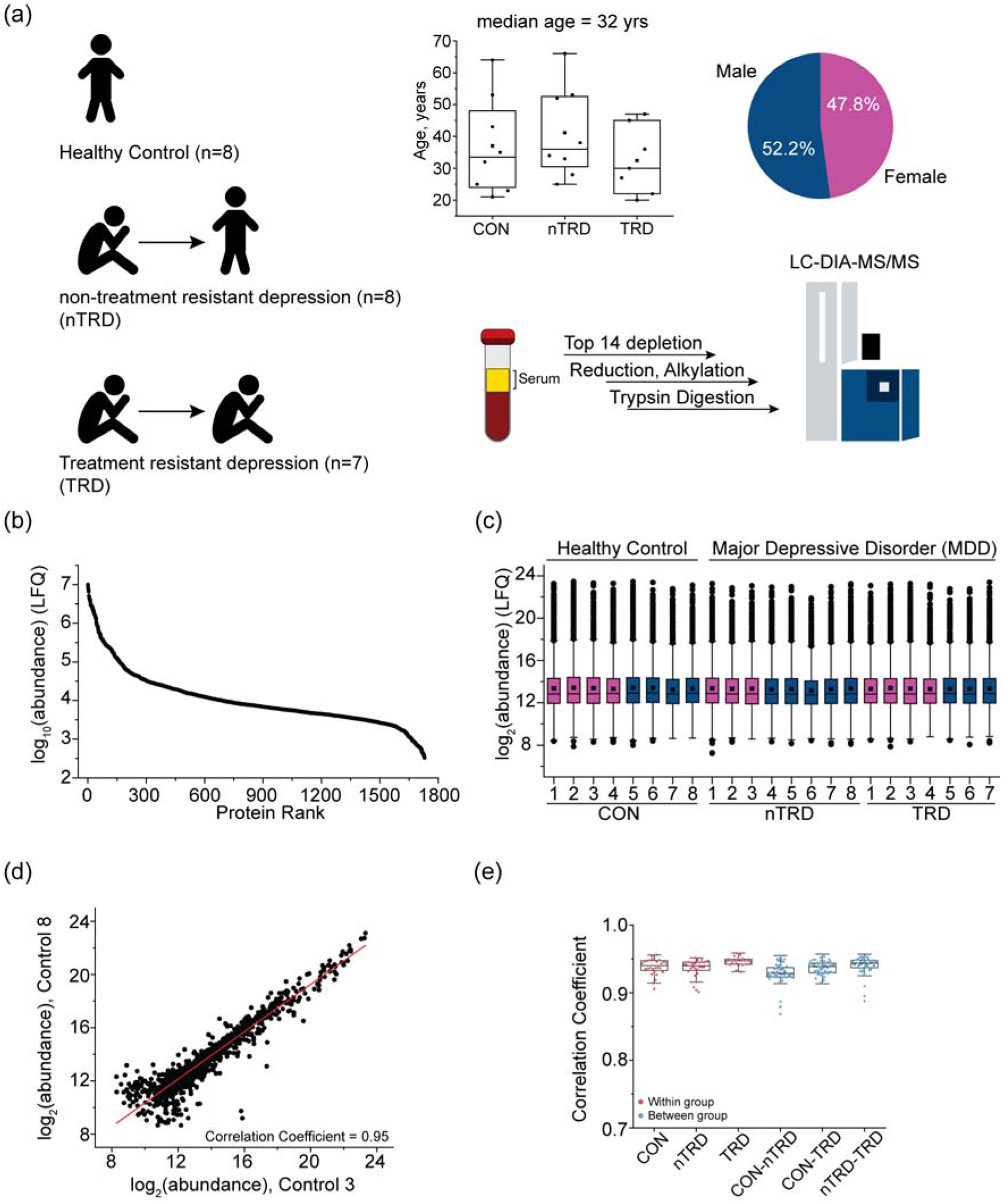
Overview of the study cohort, serum proteomics workflow, and data quality. **(a)** Study design and cohort characteristics. Serum samples were collected from healthy controls (CON, *n* = 8), non-treatment-resistant depression (nTRD, *n* = 8), and treatment-resistant depression (TRD, *n* = 7) participants. Age distribution across groups is shown by boxplots (median age = 32 years), and sex distribution is shown as percentages. Serum samples underwent depletion of the top 14 high-abundance proteins, followed by reduction, alkylation, tryptic digestion, and LC-DIA-MS/MS analysis. **(b)** Ranked distribution of serum protein abundances, demonstrating broad dynamic range and proteome depth across samples. **(c)** Distribution of log_2_-transformed protein abundances across individual serum samples in each group, indicating comparable global abundance profiles after normalization. **(d)** Representative pairwise scatter plot of log_2_-transformed protein abundances between two control samples, illustrating high technical and biological reproducibility (Pearson correlation coefficient shown). **(e)** Summary of pairwise Pearson correlation coefficients within diagnostic groups and between groups, demonstrating consistently high reproducibility across serum proteomes and comparable similarity between groups.

### Covariate-adjusted serum proteomics reveals group-associated separation

Serum protein abundances exhibited measurable associations with age and sex, exemplified by APOA1, the abundance of which were positively associated with both variables (**Fig. 2a, top**). Adjustment using a linear model incorporating age and sex as covariates effectively attenuated confounding effects. Inter-individual biological variation, was preserved as exemplified by examining the per-patient log_2_-scaled protein abundances of the residuals (**Fig. 2a, bottom**). Following covariate adjustment, PCA of the serum proteome revealed separation of CON from both depression groups along PC1 (19.77% variance) (**Fig. 2b**). nTRD and TRD occupied partially overlapping regions of the PCA space, suggesting shared proteome signatures. Gene ontology analysis of the top 25 proteins contributing to PC1 identified enrichment for extracellular exosome-associated proteins (GO CC: 0070062; -log10(FDR) = 3.88), suggesting the principal axis of variation reflects circulating extracellular protein biology in MDD relative to CON (**Fig. 2c**).

**Figure 2.**
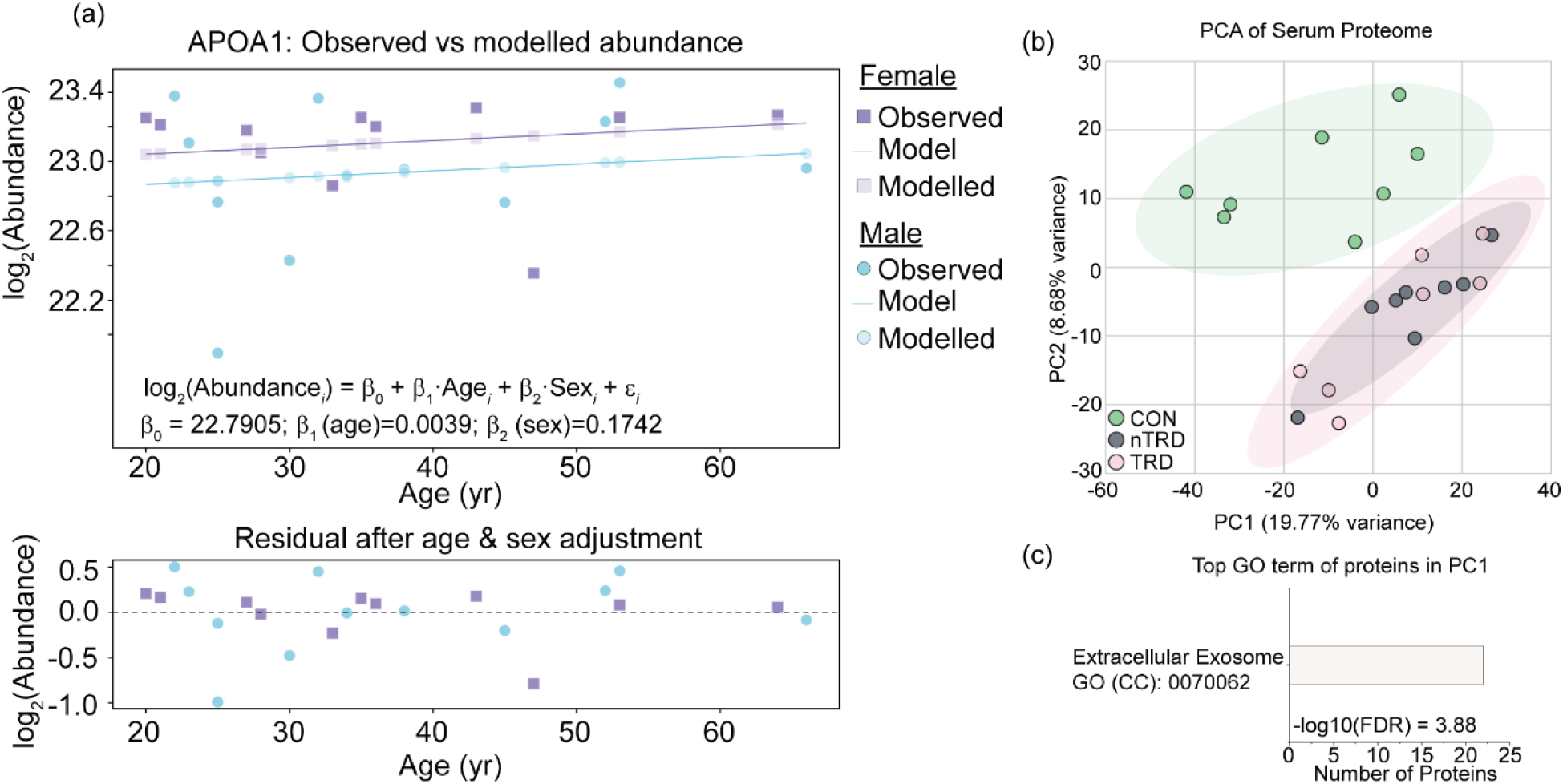
Accounting for demographic covariates and global variance structure in the serum proteome. (a) Representative example of covariate modeling for APOA1 abundance. Observed log_2_-transformed serum protein abundances are shown as a function of age, stratified by sex. A linear model incorporating age and sex effects was fitted, and model-predicted values are overlaid. Residuals after age and sex adjustment are shown below, demonstrating effective removal of demographic effects while preserving inter-individual variation. (b) Principal component analysis (PCA) of the covariate-adjusted serum proteome reveals separation between CON, nTRD, and TRD groups along the first two principal components. Shaded ellipses indicate 95% confidence interval (CI). (c) Gene ontology (GO) cellular component enrichment of the top 25 proteins contributing to PC1 highlights extracellular exosome-associated proteins, indicating that the dominant axis of variance in the serum proteome reflects circulating extracellular protein biology.

### Differential abundance and pathway enrichment in the serum proteome

Differential analysis of the covariate-adjusted serum proteome identified 56 significantly altered proteins in nTRD relative to CON (11 increased, 45 decreased; **Fig. 3a**). Proteins that were found to be downregulated included the synaptic organizer CBLN4, the monocyte/macrophage marker CD163, the extracellular matrix degrader cathepsin S (CTSS), and Ficolon-2 (FCN2), among others. Gene ontology enrichment analyses of the downregulated proteins revealed they are largely converge on terms related to immune and inflammatory processes, including positive regulation of immune responses and signalling, alongside complement activation (**Fig. 3b**). Consistently, GSEA confirmed negative enrichment of immune and complement-related pathways in nTRD, alongside positive enrichment of oxidative phosphorylation, translation, and nervous system development pathways (**Fig. 3c**). In TRD relative to CON, 26 proteins were significantly altered (6 increased, 20 decreased; **Fig. 2d**). Among many, the downregulated proteins included the immunomodulatory HLA-A, the G protein-activated inward rectifier potassium channel 3 (GIRK3), the mitochondrial NAD-dependent malic enzyme ME2, the pro-inflammatory granulin (GRN), and CTSS. Gene ontology analysis confirmed the set of downregulated genes were similarly enriched for adaptive immune and immunoglobulin-related processes (**Fig. 3e**). GSEA revealed a broadly comparable signature, with negative enrichment of immune and complement cascades and positive enrichment of oxidative phosphorylation and fatty acid metabolism (**Fig. 3f**).

**Figure 3.**
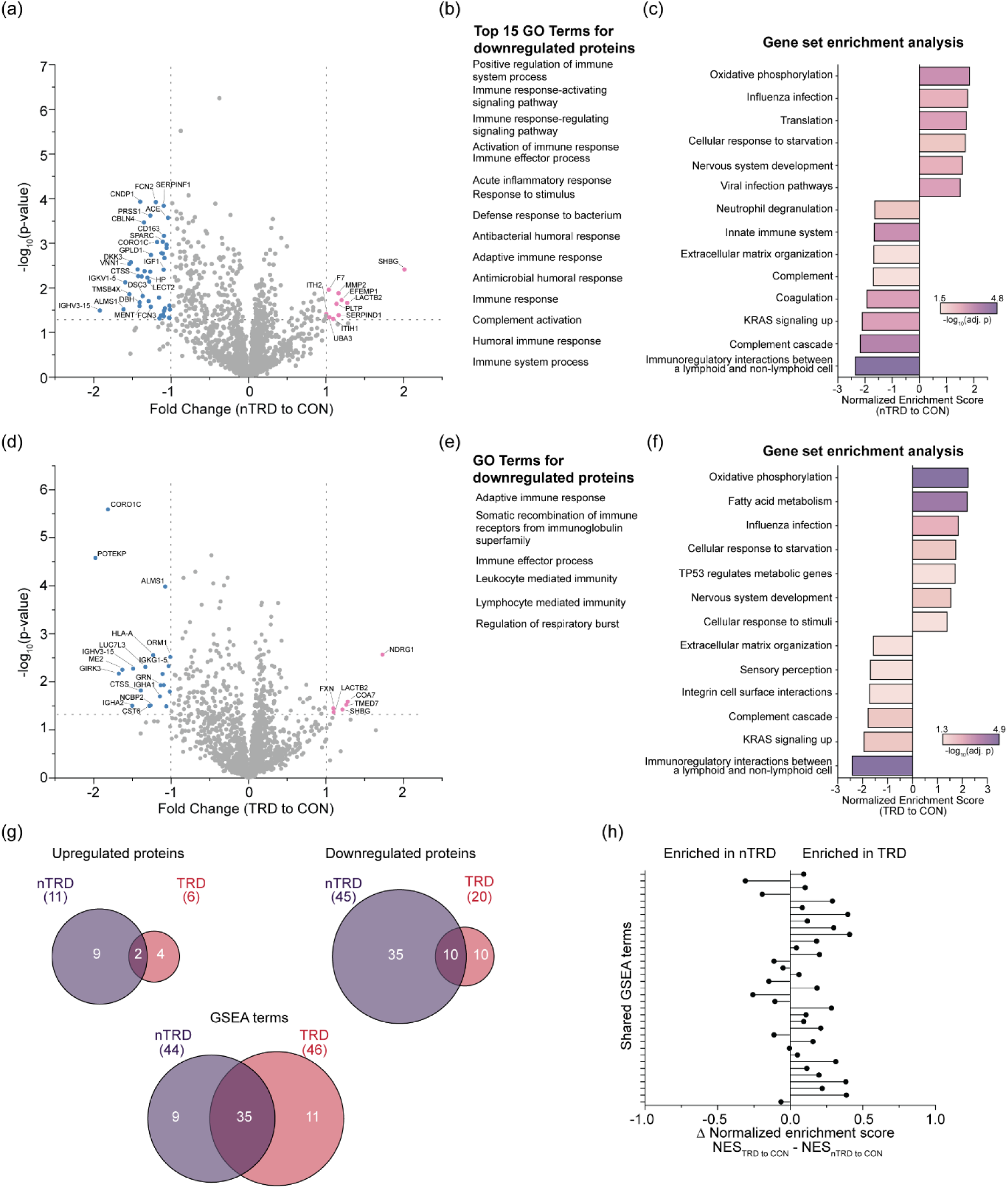
Differential protein abundance and pathway-level alterations in serum from nTRD and TRD relative to CON. (a) Volcano plot showing differentially abundant serum proteins in nTRD versus CON. Proteins significantly decreased or increased in nTRD are highlighted. (b) Gene ontology (GO) biological process enrichment of downregulated proteins in nTRD relative to controls, highlighting immune and inflammatory response pathways. (c) Gene set enrichment analysis (GSEA) for nTRD versus CON, showing normalized enrichment scores for significantly enriched pathways. (d) Volcano plot showing differentially abundant serum proteins in TRD versus CON. (e) GO biological process enrichment of downregulated proteins in TRD relative to controls, indicating altered immune and inflammatory signaling. (f) GSEA for TRD versus CON, illustrating pathway-level differences distinct from those observed in nTRD. (g) Venn diagrams summarizing overlap and divergence of significantly upregulated proteins, downregulated proteins, and enriched GSEA terms between nTRD and TRD. (h) Comparison of shared GSEA terms between nTRD and TRD, plotted as the difference in normalized enrichment scores ΔNES; few pathways are enriched in TRD or nTRD.

Direct comparison of differential protein sets revealed limited overlap between groups: 10 of 45 downregulated proteins in nTRD were also decreased in TRD, and only 2 upregulated proteins were shared (**Fig. 3g**). In contrast, pathway-level concordance was substantial: 35 of 44 significantly enriched terms in nTRD overlapped with 35 of 46 enriched terms in TRD. ΔNES analysis of shared pathways showed minimal directional divergence between groups (**Fig. 3h**), indicating that serum proteomic alterations largely reflect shared systemic biology across depressive subtypes rather than treatment-resistance-specific effects.

### EV proteome profiling reveals limited group-level divergence

The predominance of exosome-associated proteins along the primary axis of serum proteome variance prompted direct profiling of circulating EVs. EVs were isolated from serum by MagNet-based enrichment, and analysed by DIA-PASEF (**Fig. 4a**). Approximately 530 proteins were quantified across EV samples (**Fig. 4b**), including 22 annotated as brain-enriched, consistent with the capacity of circulating EVs to carry CNS-associated protein cargo. We carried out gene ontology analyses on these 22 proteins, and observed that eight of the top 10 enriched terms were heavily involved with CNS processes (axon development, neuron projection development, neuron differentiation, regulation of synapse organization, regulation of synaptic plasticity, nervous system development, NMDA glutamate receptor clustering, regulation of postsynapse organization) further highlighting the capacity for EVs to carry signatures from the brain (**Fig. 4c**). Within-group and between-group Pearson correlations were consistently high (∼0.92–0.95), confirming robust reproducibility (**Fig. 4d**). As in serum, age and sex effects were modelled and removed prior to downstream analysis, illustrated by adjustment of the canonical EV marker CD9 (**Fig. 4e**).

**Figure 4.**
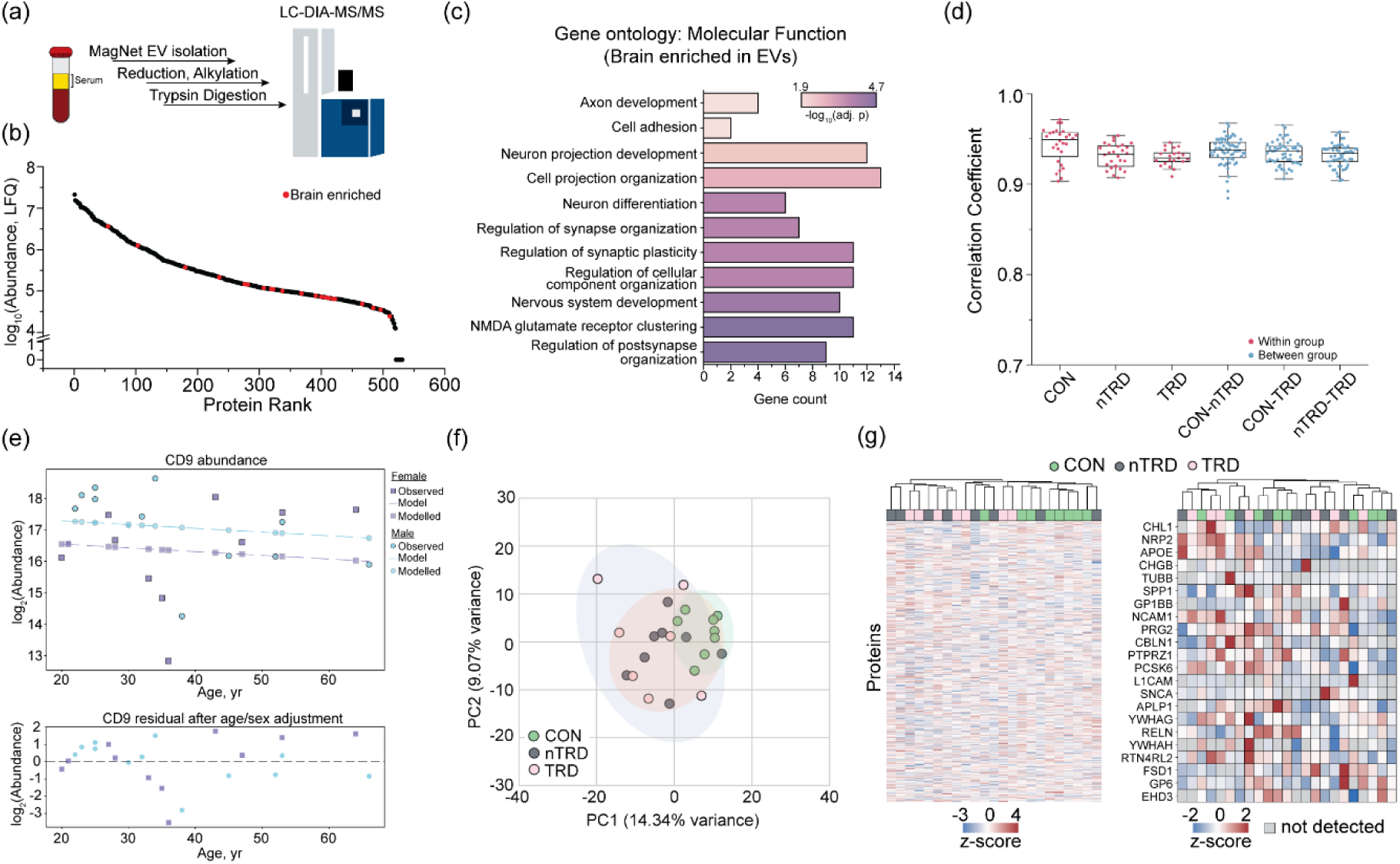
Isolation, characterization, and global structure of serum-derived extracellular vesicle (EV) proteomes across the cohort. (a) Overview of the EV proteomics workflow. EVs were isolated from serum using MagNet-based enrichment, followed by protein reduction, alkylation, tryptic digestion, and LC-DIA-MS/MS analysis. (b) Ranked distribution of EV protein abundances (log_10_ LFQ), demonstrating the dynamic range and depth of the EV proteome. Proteins annotated as brain-enriched are highlighted in red, indicating the presence of central nervous system–associated proteins within circulating EVs. (c) Gene ontology enrichment analysis (PANTHER, Fisher’s exact test, FDR-corrected) of biological process terms associated with the 26 brain-enriched proteins detected in serum-derived EVs. Terms are coloured by significance (-log_10_ adjusted p-value). Enriched terms span synaptic organization and plasticity, axon development, cell adhesion, and neuron projection development, consistent with the CNS-association of these proteins. (d) Pairwise Pearson correlation coefficients summarizing within-group and between-group reproducibility of EV proteomes. High correlation coefficients across all comparisons indicate robust analytical reproducibility and comparable global similarity between diagnostic groups. (e) Representative example of covariate modeling for the EV marker CD9. Observed log_2_-transformed CD9 abundances are shown as a function of age and sex, with fitted model predictions overlaid. Residuals after age and sex adjustment are shown below, illustrating effective removal of demographic effects while preserving biological variability. (f) PCA of covariate-adjusted EV proteomes. Samples from CON, nTRD, and TRD show substantial overlap along the first two principal components, indicating limited global separation and pronounced inter-individual heterogeneity in EV protein cargo. Shaded ellipses denote the 95% confidence intervals. (g) Unsupervised hierarchical clustering of covariate-adjusted EV proteomes (left) and brain-enriched protein (right) visualized as a z-score–scaled heatmap. In both cases, no strong group-wise clustering is observed.

In contrast to serum, PCA of covariate-adjusted EV proteomes revealed no clear group-level separation: CON, nTRD, and TRD substantially overlap along PC1 (14.34% variance) and PC2 (9.07% variance), with pronounced inter-individual heterogeneity (**Fig. 4f**). Hierarchical clustering of z-score-scaled EV protein abundances similarly did not reveal strong treatment group segregation from CON subjects. Group-wise separation was not observe following hierarchical clustering of the 22 brain enriched proteins found in the EV proteome. (**Fig. 4g**). This suggests that large differences in protein abundances in EVs are not distinguishers of treatment group in our psychiatric cohort.

### Distinct EV proteomes reveal divergent pathway-level signatures in nTRD and TRD

Despite limited global separation by PCA, differential analysis of EV proteomes revealed distinct protein- and pathway-level changes in nTRD and TRD relative to CON (**Fig. 5**). In nTRD EVs, 45 proteins were significantly altered (23 increased, 22 decreased; **Fig. 5a**). Upregulated proteins included components involved in translational elongation (EEF2), protein folding and quality control (TCP1, PSMA2), mitochondrial energy metabolism (SLC25A3), and antioxidant defence (PON3), alongside the synaptic scaffold protein YWHAH and the cell adhesion molecule CADM1. Interestingly, proteins involved with extracellular matrix remodelling (MMP19, DCN, LTBP2, TNN) were also upregulated, consistent with previous reports of metalloprotease-mediated extracellular matrix degradation in the nucleus accumbens in a mouse model of chronic stress.^35^ Downregulated proteins included cytoskeletal components (TUBA4A, MYL12B), the endosomal recycling GTPase EHD3, the neuroendocrine dense-core vesicle protein chromogranin-B (CHGB), the mitochondrial chaperone HSPD1, the glucose transporter GLUT1 (SLC2A1), and innate immune proteins including ficolin-2 (FCN2) and lactoferrin (LTF). While these proteins did not yield significantly enriched GO terms, GSEA of EV proteomes in nTRD relative to CON identified positive enrichment of oxidative phosphorylation, mitochondrial protein degradation, aerobic respiration, and MYC targets alongside negative enrichment of vesicle-mediated transport, complement cascade, and intracellular signaling pathways (**Fig. 5b**).

**Figure 5.**
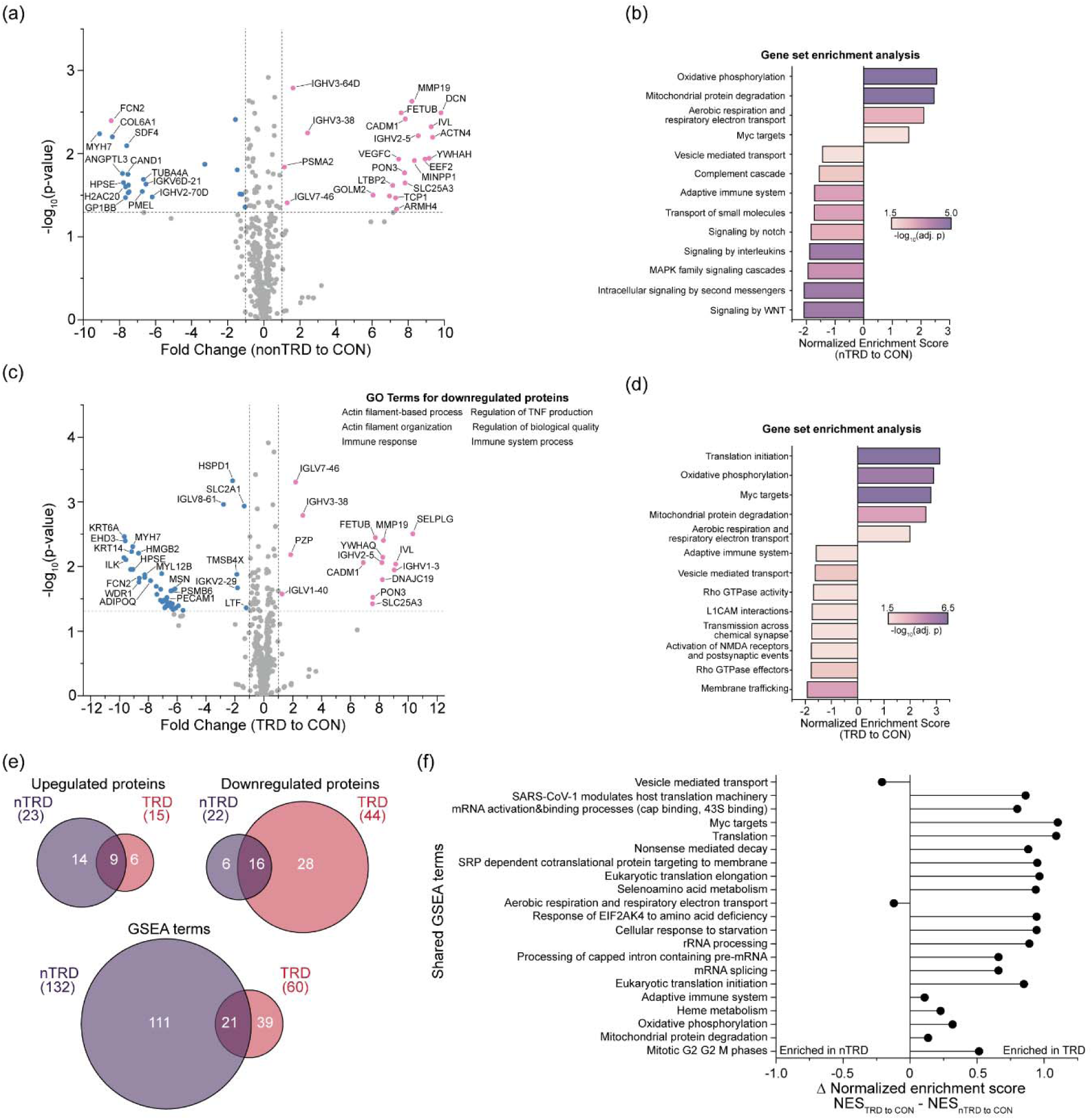
Differential abundance and pathway-level enrichment of serum-derived EV proteomes in nTRD and TRD compared with CON. (a) Volcano plot showing differentially abundant EV proteins in nTRD versus CON. Proteins significantly increased or decreased in nTRD are highlighted. (b) GSEA of EV proteins in nTRD versus CON, displaying normalized enrichment scores for selected significantly enriched pathways. (c) Volcano plot showing differentially abundant EV proteins in TRD versus CON. (d) GSEA of EV proteins in TRD versus CON, highlighting pathway-level changes distinct from those observed in nTRD. (e) Venn diagrams summarizing overlap and divergence of significantly upregulated proteins, downregulated proteins, and enriched GSEA terms between nTRD and TRD. (f) Comparison of shared GSEA terms between nTRD and TRD expressed as the difference in normalized enrichment scores (ΔNES), illustrating pathways are relatively more enriched in TRD despite shared statistical significance.

Relative to CON, 59 proteins were significantly altered in TRD EVs (15 increased, 44 decreased; **Fig. 5c**). Relative to CON, 59 proteins were significantly altered in TRD EVs (15 increased, 44 decreased; Fig. 5c). Upregulated proteins included mitochondrial bioenergetic and import machinery components (SLC25A3, DNAJC19), the antioxidant enzyme PON3, the ECM metalloproteinase MMP19, synaptic scaffolding proteins (YWHAQ, CADM1), and secreted protease inhibitors (PZP, FETUB). Downregulated proteins spanned multiple functionally coherent groups: actin cytoskeletal and motor proteins (MYL12B, MYH7, MSN, TAGLN2, TMSB4X, WDR1); the microtubule subunit TUBA4A; synaptic and dendritic proteins including drebrin-1 (DBN1) and, the sheddase ADAM10; the endosomal recycling GTPase EHD3; the neuroendocrine secretory protein chromogranin-B (CHGB); the mitochondrial chaperone HSPD1; metabolic regulators (SLC2A1, ADIPOQ, CD36); and innate immune proteins (FCN2, LTF, MPO). Downregulated proteins were significantly enriched for terms associated with actin filament organization, regulation of TNF production, and immune regulated processes (**Fig. 5c inset**). GSEA demonstrated a polarized pathway signature: strong positive enrichment of translation initiation, oxidative phosphorylation, MYC targets, mitochondrial protein degradation, and aerobic respiration was accompanied by pronounced negative enrichment of pathways related to membrane trafficking, synaptic transmission, activation of NMDA receptors, L1CAM interactions, and Rho GTPase signaling (**Fig. 5d**).

Direct comparison showed nTRD and TRD shared 9 upregulated and 16 downregulated proteins, with TRD additionally exhibiting a substantially larger number of uniquely dysregulated proteins (**Fig. 5e**). At the pathway level, 132 GSEA terms were significant in nTRD and 60 in TRD, with 21 shared between groups. ΔNES analysis of shared pathways indicated enhanced enrichment of metabolic, translational, and RNA processing pathways – including mitotic G2/G2M phases, oxidative phosphorylation, heme metabolism, and translation initiation – in TRD relative to nTRD (positive ΔNES), whereas vesicle-mediated transport showed relatively greater enrichment in nTRD (negative ΔNES; **Fig. 5f**). These results suggest selective amplification of bioenergetic and translational programs within TRD-derived EVs, accompanied by relative depletion of synaptic and membrane-trafficking signatures, patterns that were not captured by bulk serum proteomics.

## Discussion

In this exploratory study, we profiled paired serum and serum-derived EV proteomes from healthy controls and individuals with nTRD and TRD using DIA-PASEF mass spectrometry. Our principal finding is that, while bulk serum proteomes exhibited detectable global separation between depression groups and controls, these changes are largely shared across nTRD and TRD and do not resolve treatment-resistance-specific biology. By contrast, despite limited global separation, EV proteomes had TRD-specific pathway disruptions, such as unexpected downregulation of immune pathways, and amplified mitochondrial metabolism, oxidative phosphorylation, and translational programs.^23,24^ Consistent with the paradigm that stress imposes persistent changes to molecular and cellular circuitry in the brain,^35,36^ the TRD group also showed depletion of synaptic signaling, membrane trafficking, and cytoskeletal pathways which were specifically captured in the EV proteomes.

Despite our unexpected observation that serum proteomes in both nTRD and TRD share downregulated immune and inflammatory pathway changes (relative to CON), the disruption is consistent with the established role of neuroinflammation in MDD pathophysiology. Reduced immune effector and complement proteins in serum may reflect altered innate immune regulation that is common to depression regardless of treatment responsiveness. However, the near-complete overlap of significant GSEA terms between nTRD and TRD in serum, and the minimal ΔNES divergence between groups (**Fig. 3h**), indicate that bulk serum proteomics provides insufficient resolution to stratify depressive subtypes at the pathway level in this cohort.

EV proteomes capture a more selective snapshot of intercellular signaling.^18^ Our data raises the possibility that, even in this small preliminary cohort, TRD can be distinguished from nTRD at the pathway level. This possibility is consistent with recent plasma EV proteomic studies which resolved clinically and biologically distinct subtypes of Parkinson’s Disease^37^ and Alzheimer’s Disease^38^ which were indistinguishable at bulk plasma proteome level. The enrichment of mitochondrial bioenergetic and protein quality-control programs in TRD EVs is of particular interest. Mitochondrial dysfunction and oxidative stress have been implicated in MDD and TRD by both transcriptomics^14,39^ and metabolomics^40^. Our parallel proteomic findings extend these observations to circulating EV cargo. Despite the small size of the cohort, the concomitant enrichment of MYC-regulated programs and translational initiation pathways suggests broad upregulation of anabolic and biosynthetic programs within EV-producing cells in TRD, which provides the exciting possibility that brain-derived EVs can carry molecular signatures from the CNS into circulation. Together, these findings support a model in which altered cellular energy metabolism and proteostasis are reflected in the EV component of individuals with TRD, even in this modest pilot cohort

While individual protein abundances in EVs did not appear to drive separation between clinical groups, it is striking that we captured CNS-enriched proteins found in EVs showed pathway-level changes (synaptic transmission, NMDA receptor function, and L1CAM-mediated cell adhesion) in TRD relative to CON. These strong signatures were revealed in our small cohort, and are consistent with synaptic dysfunction and impaired neural connectivity as candidate mechanisms of antidepressant resistance, and align with preclinical evidence that EV-mediated synaptic communication is disrupted in models of chronic stress and mood disorders. It is possible that the loss of membrane trafficking and cytoskeletal signatures from TRD EVs may further reflect impaired vesicle biogenesis or sorting in neurons and glia, although cell-type specific EV isolation would be required to assign these cargo changes definitively to cellular origins.

While this exploratory, cross-sectional analysis of a small cohort (n = 23) establishes proof-of-concept for the utility of deep, untargeted proteomics as a viable approach for understanding proteome signatures in depression subgroups, several limitations of the present study warrant consideration. The assignment of nTRD and TRD status was based on a common – but not universal – definition in the field of failure to response to two or more antidepressant treatments in the current episode. Further studies using alternative TRD definitions will be needed to confirm the generalizability of these findings. Moreover, while EV proteomes are enriched for brain-expressed proteins, EV cargo is contributed by many cell types and the cellular origin of specific pathway signatures cannot be inferred without cell-type-specific isolation methods. With continued improvements to enrichment strategies, focused on isolating CNS-derived EVs from serum and cerebrospinal fluid^26^, direct observation of brain-derived EV cargo and signatures altered across depressive subtypes and treatment response categories will become increasingly tractable, providing more direct windows into maladaptive cellular processes in affective disorders and bringing the field closer to proteomic stratification of antidepressant treatment failure. Collectively, such advances may not only improve patient stratification but shed light on the intercellular signalling disruptions that drive MDD pathophysiology more broadly.

In summary, this study demonstrates that serum-derived EV proteomics provides a complementary and more informative window into depression biology than bulk serum proteomics. The treatment-resistance-specific pathway signature identified in TRD EVs — encompassing altered bioenergetics, translational dysregulation, and synaptic depletion — highlights EV-based profiling as a promising approach for the discovery of biomarkers and mechanistic insights into antidepressant resistance.

## Data Availability

All mass spectrometry data have been deposited to the ProteomeXchange Consortium via the PRIDE^41^ partner repository with the dataset identifier PXDXXXXXX.

## Acknowledgements

Work on “A trans-omic platform to define molecular interactions underlying anhedonia at the blood–brain interface” (C.V.R. and T.J.E.) and “Defining Liquid Biosignatures of Treatment-Resistant Anhedonic Depression in Humans” (J.W.M. and S.J.R.) are supported by Wellcome Leap as part of the Multi-Channel Psych Program. Work in the C.V.R. laboratory is also supported by a Louis Jeantet Award, a Wellcome Trust Award (221795/Z/20/Z), and a European Research Council (ERC) Advanced Grant (EP/Y029259/1). J.W.M also acknowledges support provided by the Ehrenkranz Center for Human Resilience at the Icahn School of Medicine at Mount Sinai and by a generous support from the Gottesman Foundation and Great Hill. W.B.S. acknowledges funding from the UKRI Future Leaders Fellowship (MR/V02213X/1) and the Royal Society (RG/R1/241103). S.A.B. acknowledges funding from the Wellcome Trust Early Career Award (312750/Z/24/Z) and a Dobson Junior Research Fellowship in Molecular Biophysics from Lady Margaret Hall.

